# dsRNAEngineer: a web-based tool of comprehensive dsRNA design for pest control

**DOI:** 10.1101/2024.07.22.604585

**Authors:** Yang Chen, Yufei Shi, Ziguo Wang, Xin An, Siyu Wei, Christos Andronis, John Vontas, Jin-jun Wang, Jinzhi Niu

**Affiliations:** Key Laboratory of Entomology and Pest Control Engineering, College of Plant Protection, Southwest University, Chongqing, China; Key Laboratory of Agricultural Biosafety and Green Production of Upper Yangtze River (Ministry of Education), Southwest University, Chongqing, China; Institute Molecular Biology and Biotechnology, Foundation for Research and Technology, Heraklion, Crete, Greece; Agricultural University of Athens, Athens, Greece

## Abstract

RNA interference (RNAi) is a form of gene silencing triggered by double-stranded RNA (dsRNA) that operates in all eukaryotic organisms, including insects, mites, nematodes and fungi. In the last two decades, many dsRNAs have been synthesized to silence the target genes for exploration of the underlying function in these pests. Some of them are lethal to pests or inhibit the growth of the pest population, leading to a new concept of pesticides. The generation of these environment-friendly pesticides requires precise *in silico* design of dsRNA molecules that are specific to target pests but do not target non-pest organisms. Current efforts for dsRNA design are mostly focused on gene sequence level, lacking comprehensive analysis of RNAi-based mode-of-action at the whole transcriptome level of given species, causing low efficiency and imprecise dsRNA target exploration. To address these limitations, we have created dsRNAEngineer, a publicly available online tool that allows a comprehensive and rational dsRNA design, incorporating hundreds of pests and non-pests transcriptomes. Developed functionalities include screen-target (screen conserved genes for co-targets of various pest species), on-target, off-target, and multi-target to generate optimal dsRNA for precise pest control. dsRNAEngineer is available at https://dsrna-engineer.cn/.

**Graphical abstract:** 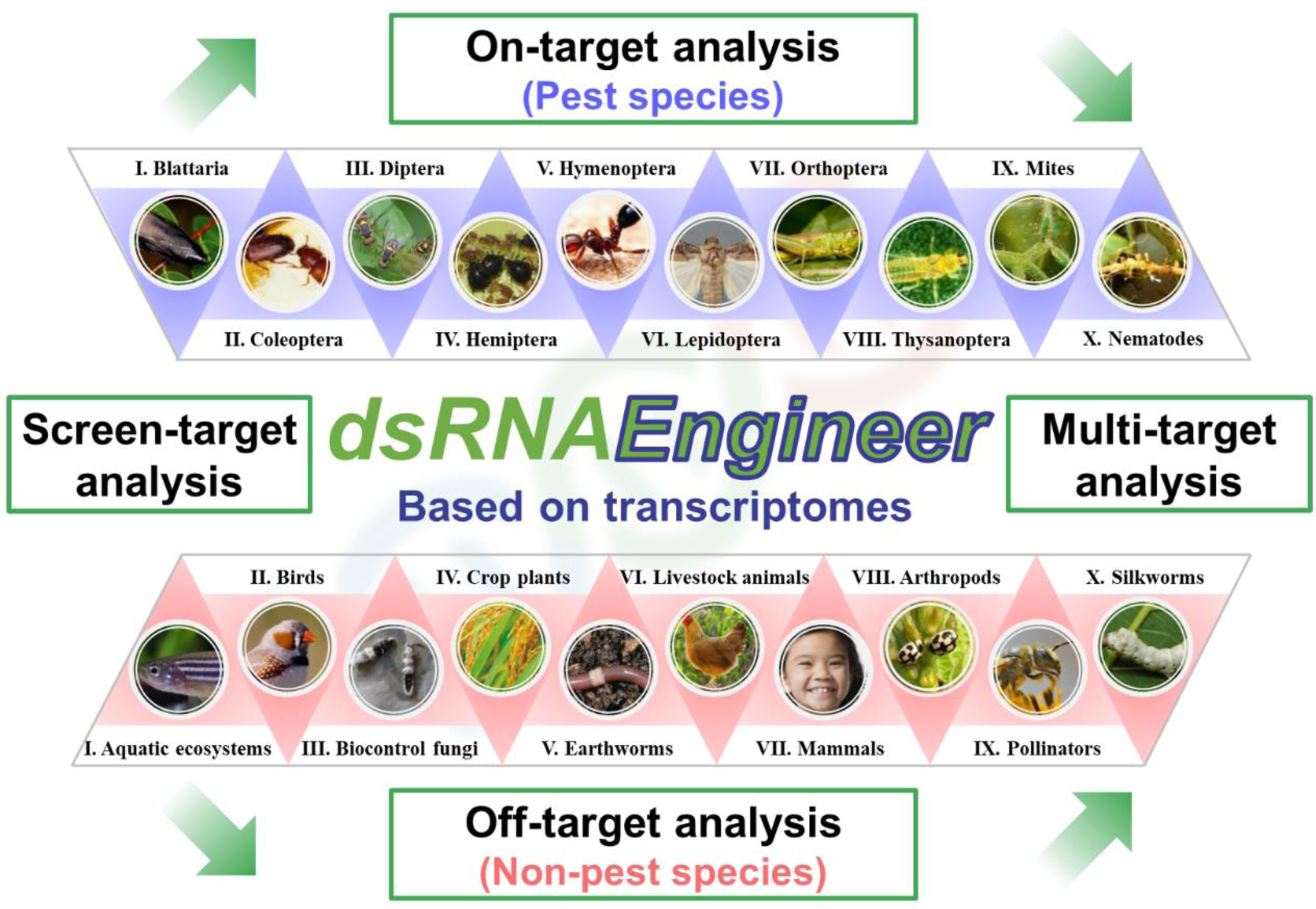

## Introduction

The mechanism of RNA interference (RNAi) was first reported in 1998 in the nematode *Caenorhabditis elegans* (1). This discovery was awarded the 2006 Nobel Prize in Physiology or Medicine. This mechanism has since been discovered in all eukaryotic organisms including insects, mites, nematodes and fungi, organisms that damage crops and hence, pose a great threat to global food security (2). To date, the threat of pests has been mostly addressed with chemical/synthetic pesticides which pose a considerable harm to the environment (3). An alternative strategy to combat pests is to take advantage of the inherent RNAi mechanisms that act in a sequence-specific manner for gene silencing, and can be triggered by exogenously synthesized dsRNA. Triggering RNAi via the usage of dsRNAs has been intensively explored in various pest and non-pest species to investigate gene function (4,5). Many dsRNAs have been synthesized to silence target genes for exploration of functions in various pests, and some of these dsRNA can lead pests to lethality or inhibit the growth of the pest population, providing a new generation of pesticides (6,7). Recently, two RNAi-based pest control products were registered, one is the transgenic maize (MON 87411) engineered to express dsRNA in targeting *sucrose non-fermenting protein 7* (*Snf7*) for control of western corn rootworm (*Diabrotica virgifera virgifera*), another is a sprayable dsRNA (Ledprona) that targets *proteasome subunit beta type-5* (*PSMB5*) for control of Colorado potato beetle (*Leptinotarsa decemlineata*) (8,9).

The key to this strategy is the rational dsRNA design by specifically targeting pests (herein termed “on-target” organisms) but at the same time avoiding non-pests (termed “off-targets”). The mode of action of a dsRNA pesticide is the following: The dsRNA molecule is cut by the pest endonuclease Dicer into 20-22 bp small interfering RNAs (siRNAs), one siRNA strand is then captured by the protein Argonaute (Ago) to form the RNA-induced silencing complex (RISC). RISC scans for complementary mRNA molecules and targets them for degradation, ultimately suppressing the target gene expression. Therefore, the key of rational dsRNA design is to follow the RNAi-based mode-of-action on gene sequence, specifically checking the specificity of small RNAs (*e.g* siRNAs), to on-target organisms compared to off-target (non-pests) ones. Past studies have indicated that small RNAs ranging from 16 bp to 26 bp could also trigger a certain level of gene silencing (10–14). In addition, off-target effects at the whole transcriptome level are a significant and non-ignorable factor, resulting in the silencing of many unintended genes (15,16). Thus, a comprehensive bioinformatic analysis for on- and off-target siRNA effects at the whole transcriptome level should include both testing the 20-22 bp range of molecules (dependent on species-specific manner of Dicer), but also two additional small siRNA sizes (16 bp perfect match and 26 bp imperfect match). Apart from the necessary domain expertise, this kind of analysis often requires programming skills. Existing online tools (17–19), mainly developed nearly a decade ago, only focus on limited model organisms in optimizing dsRNA/siRNA specificity. Moreover, current tools for dsRNA design in pest control are mostly focused at the gene sequence level, lacking comprehensive analysis of RNAi-based mode-of-action in organism transcriptome level, causing low efficiency and sometimes imprecise dsRNA target exploration.

To overcome these challenges, we developed a web-based tool called dsRNAEngineer, which incorporates hundreds of pests and non-pests transcriptomes to allow a comprehensive dsRNA design for pest control. Beyond the typical on- and off-target analysis for dsRNA pesticides, we also developed screen-target functionality to explore a conserved dsRNA target sequence in a given pest group, and multi-target functionality to optimize a dsRNA to target on multiple genes in pests.

## Materials and methods

### Web server implementation

The web server of dsRNAEngineer was built using Vue.js framework (front-end, https://vuejs.org) and Python’s asynchronous framework (back-end, https://sanic.dev). The interactive network display is created with Canvas API.

### Data collection

Currently, dsRNAEngineer hosts 942 transcriptomes from the NCBI database including worldwide insect pests, mites, nematodes and fungi, and safety-assessment species such as aquatic ecosystems, birds, pollinators, silkworms, beneficial arthropods, earthworms, biocontrol fungi, crop plants, livestock animals, and mammals (updated on the 5^th^ of July, 2024). Among them, 439 RefSeq-transcriptomes (GCF) were directly retrieved from NCBI Reference Sequence Database, and others were collected from raw reads of the Sequence Read Archive database, quality-controlled by fastp (20), assembled by Trinity (21) and annotated through the NCBI Non-Redundant (NR) database by BLAST. Although the current transcriptomes cover the majority of pest species and all required safety-assessment species, we make our best to also constantly update a collection of transcriptomes from user’s feedback and from the National Center for Biotechnology Information (NCBI).

### Work pipelines

The *Start dsRNAEngineer* tab is the gateway to the dsRNA design workflow containing two pipelines (Figure 1). The first pipeline starts from user-defined pest species in screening target genes (screen-target) and follows the on-target, off-target and multi-target analysis (Figure 1A). The second pipeline starts from user-defined target genes, and follows the on-target, off-target and multi-target analysis (Figure 1B). Once the project task is submitted, the user can use the project ID to check the status and an optional email alert on project status is also provided.

**Figure 1.**
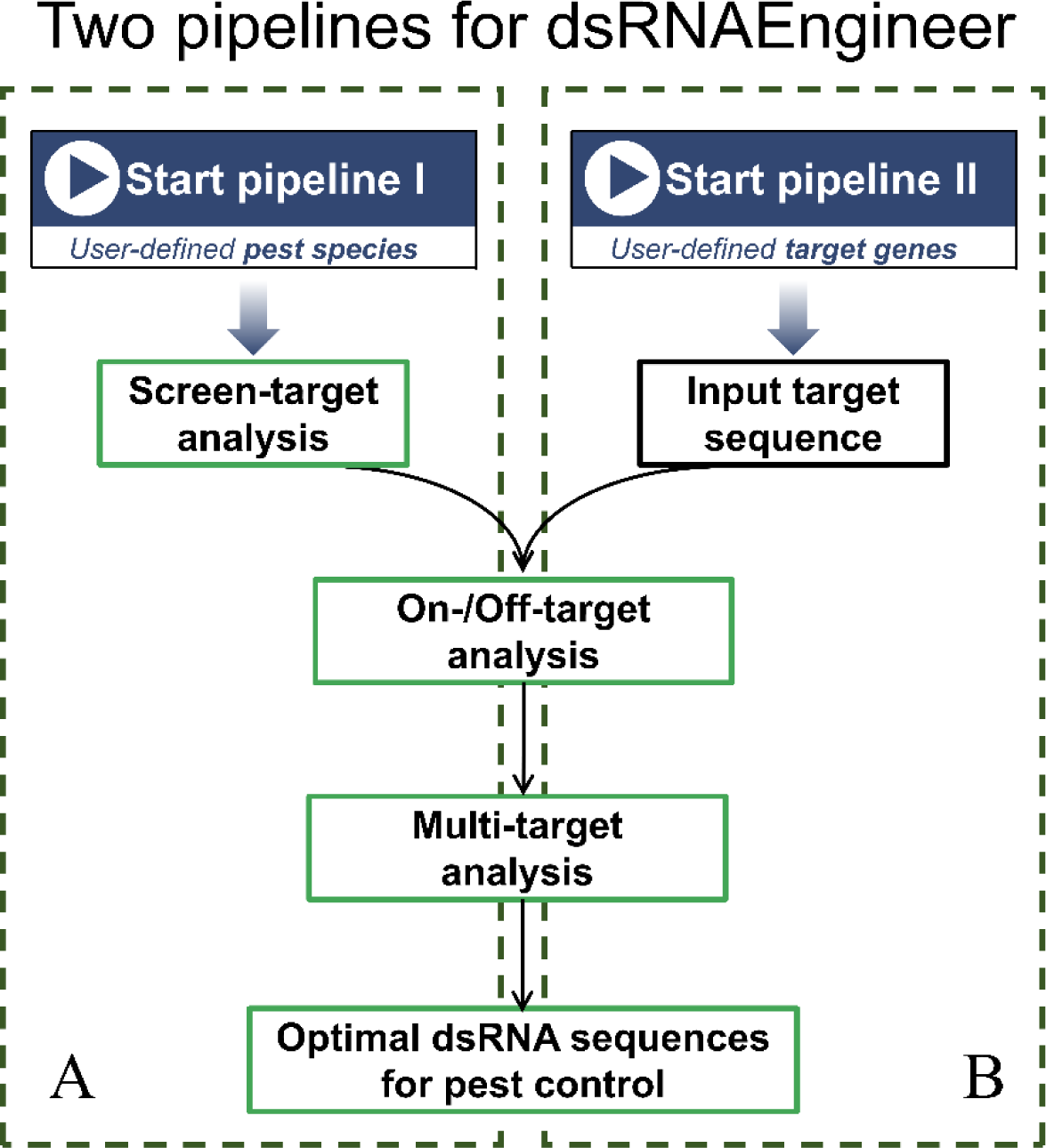
Two pipelines for dsRNAEngineer. (A) Pipeline I is suitable for user-defined pest species, performs initially a step of Screen-target to obtain target genes for the defined pest species; (B) Pipeline II is suitable for user-defined target genes through input target sequence. Following the input of either a target species or a target gene, both pipelines follow the on-/off-target analysis and multi-target analysis to generate optimal dsRNA sequences for pest control.

### Screen-target analysis: To obtain the genes that can be the candidate RNAi targets to control various pests

To start screen-target analysis (Figure 2A), a species from the major target pest list and up to four species from a list of co-target pests should be chosen to perform transcriptome level pairwise BLAST (22). After the analysis is done, the Project detail section gives the basic information of screen-analysis, and a Wayne diagram indicate the numbers of the candidate target genes in combinations of various pests. Users can view the homologous genes between two or more pests with the Wayne diagram. By default, common candidate target genes to all analyzed pests are displayed (Figure 2B). In Results section, the user can obtain the top candidate target genes by setting sequence identity, hit length, and species combination (Figure 2C). Although sequence identity among genes has been recommended to be ≥ 80% (13,23–25), it is advised to set it as high as possible to obtain the best target in targeting multiple pests. Once the candidate target genes are chosen by the *send to target* tab, these genes will be listed in target section to *proceed to on-/off- target analysis* (Figure 2D).

**Figure 2.**
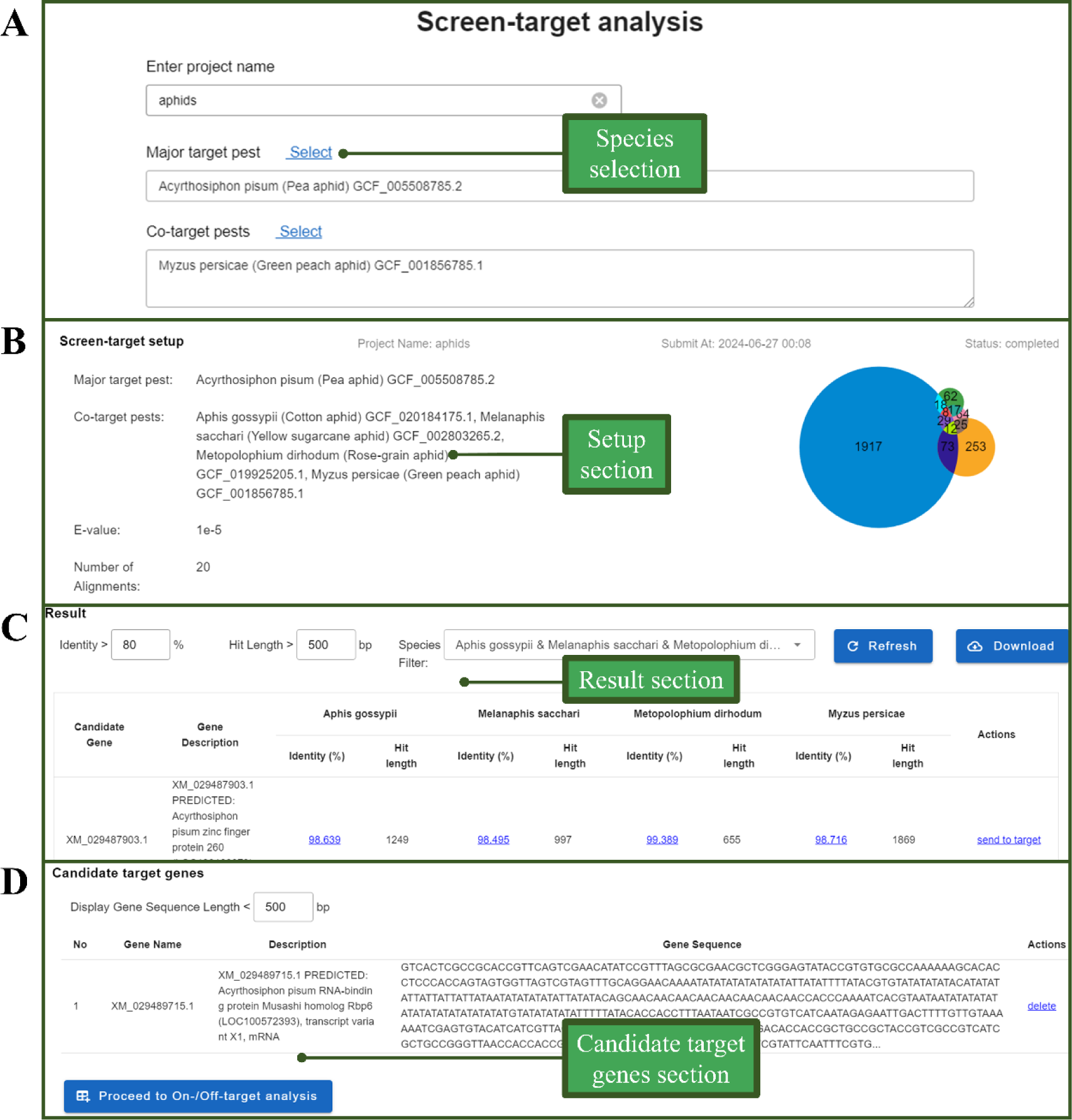
Screen-target analysis. (A) Species selection of major target pest and co- target pests. (B) Setup section: information of analysis setup and Wayne diagram for pairwise blast of genes of transcriptomes from selected pest species; (C) Results section: parameters for setting sequence identity, hit length, and species combination; (D) Candidate target genes section: genes selected for further on-/off-target analysis.

### On- and Off-target analysis: evaluate the RNAi-based on- and off- target effect to target genes for pest species and non-pest species

Together with the input target gene sequence (*Enter your target gene sequence in FASTA format*) either from the first pipeline by the user-defined pest species in screening target genes (screen-target) or from the second pipeline by the user-defined target genes, users can choose the species from both on-target list (pests) and off-target list (non-pests) for On- and Off-Target analysis (Figure 3). In addition, users can also upload FASTA files of up to 1MB in size for custom sequence On- and Off-target analysis. Two match conditions (*Default Match Model* or *Custom Options*) are provided to set the calculation model. In the Default Match Model, dsRNAEngineer performs sequential matching of the major small RNA size (the size of dsRNA sliced by Dicer-2 is predefined in a species-specific manner), and two additional small RNA sizes (16 bp perfect match and 26 bp almost perfect match with 0∼2 mismatches). In the Custom Options mode, the user can define the major small RNA size as well as two additional small RNA sizes. Then, On- and Off-Target analysis will be initiated by *On- and Off-Target analysis* tab (Supplemental Figure 1).

**Figure 3.**
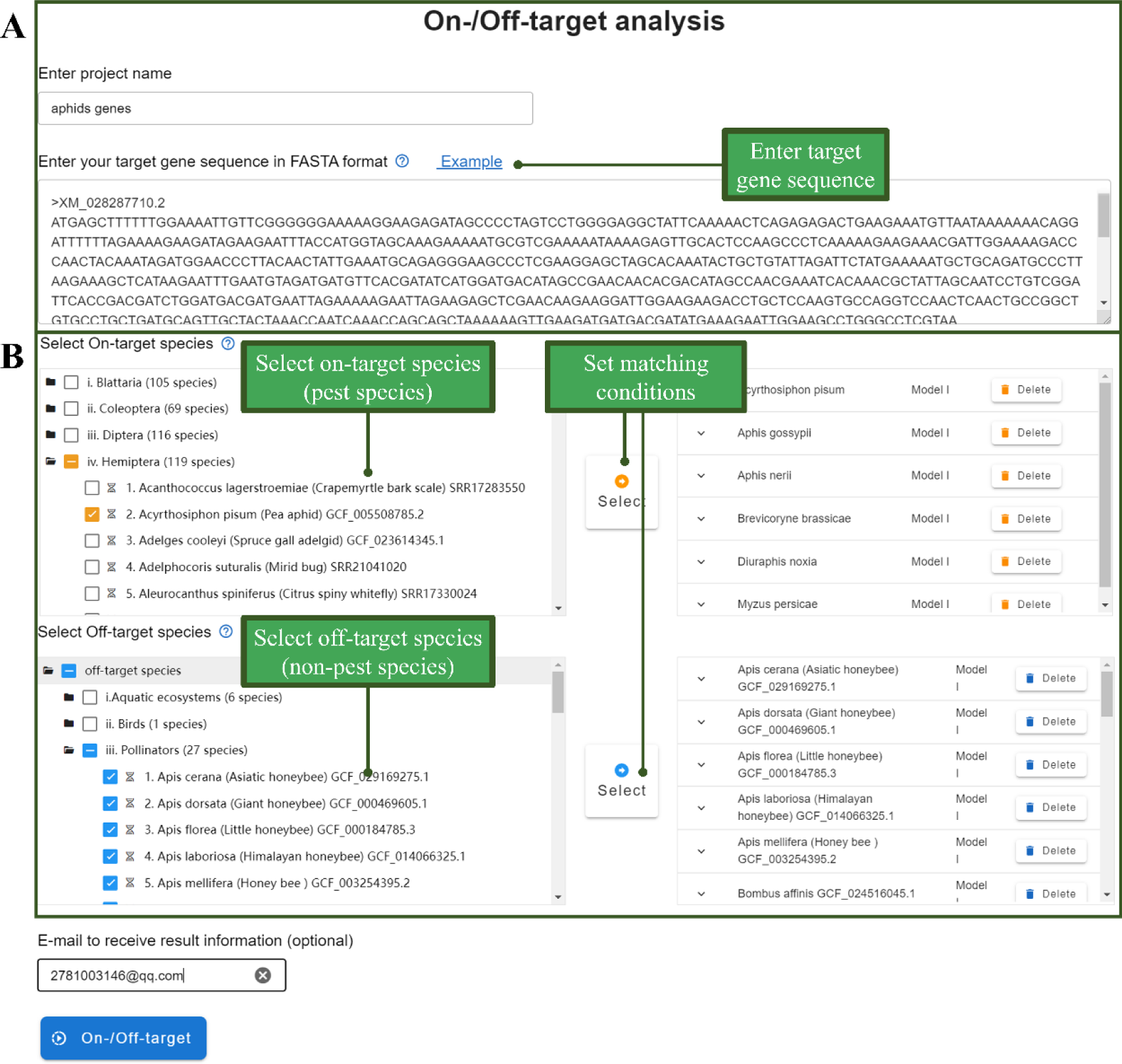
On-/Off-target analysis. (A) Input of target gene sequence either from user-defined pest species in screening target genes (screen-target) or from user-defined target genes. (B) Selection of the species from on-target list (pests) and off-target list (non-pests), and analysis model.

In the result page of On- and Off-Target analysis, the analysis setup (target gene sequences, On-target species, and Off-target species) are automatically presented on the top (Figure 4A). The analysis results are displayed with a graphical view and a table summary. In the graphical view (Figure 4B), the number of on-targets and off-targets can be visualized at different nucleic acid positions for each input target gene. The graphs can be easily visualized in different on-target species (selection from *On-target species*) and off-target species (selection from *off-target species*) and saved locally. For table summary (Figure 4C), the top candidate dsRNA fragments will be listed upon the selection of candidate dsRNA fragments (priority to pest control: on-target, or priority to biosafety: off-target), fragment size. Based on the user-selected dsRNA fragment sizes (e.g. 50-100bp up to 650-700bp in increments of 50 bp), an exhaustive sliding window analysis is performed. Finally, the top ten dsRNAs per dsRNA fragment sizes interval are recommended to the user (Supplemental Figure 2). The proposed optimal dsRNA can be chosen by selecting *send to design*. Additionally, users can also perform off-target analysis for the study of gene function in avoiding the off-target risk to non-target genes of an organism.

**Figure 4.**
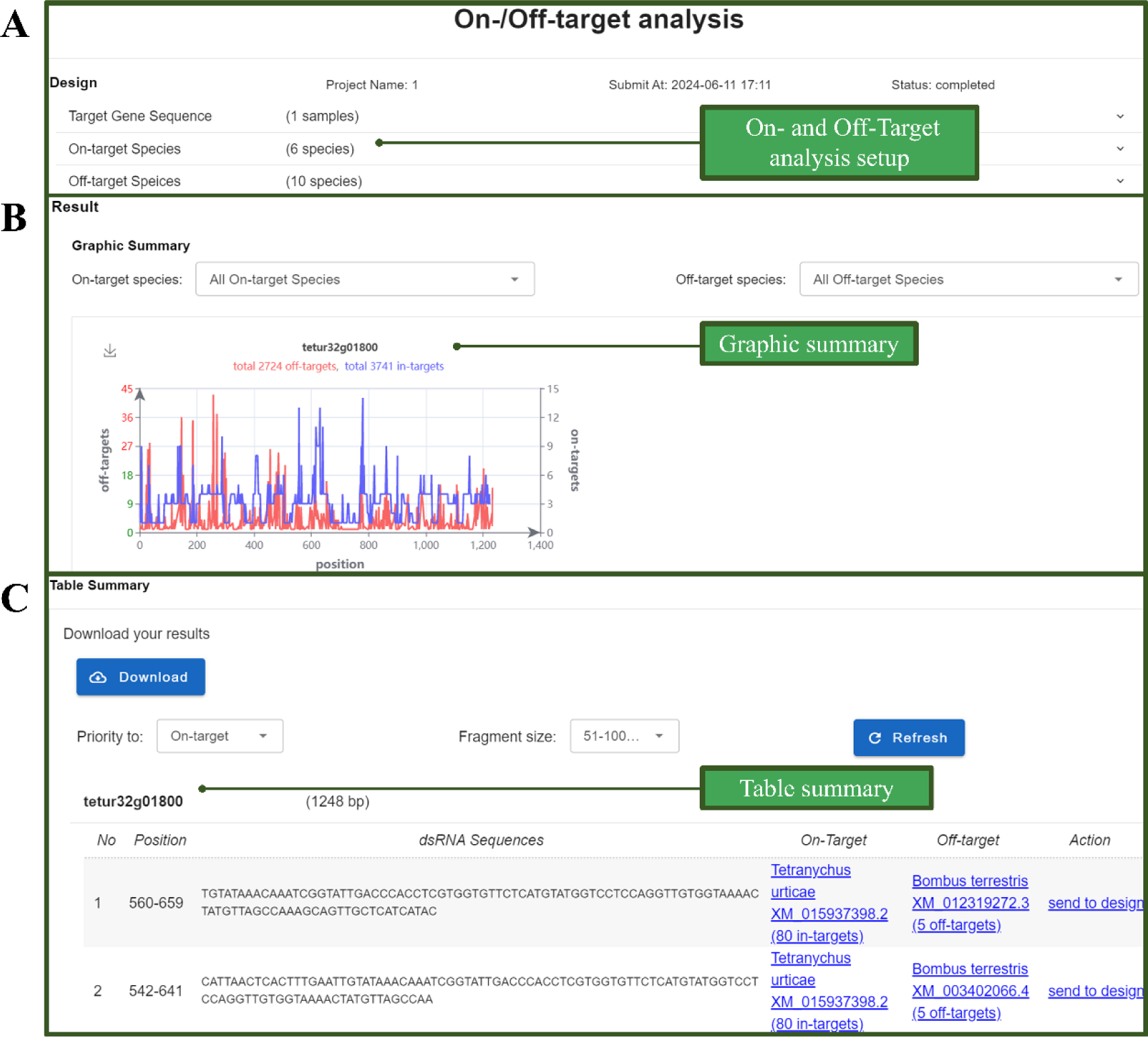
On-/Off-target analysis output. (A) Analysis setup: target gene sequences, On-target species, and Off-target species; (B) Graphic summary: the number of on- targets and off-targets can be visualized at different nucleic acid positions for each input target gene; (C) Table summary: list of top candidate dsRNA fragments.

### Multi-target analysis: design of a dsRNA fragment in targeting multiple genes

The selected dsRNA fragments will be listed in the section of *dsRNA in targeting single gene*, and these dsRNA sequences can be used as templates for further dsRNA synthesis (Figure 5A). Alternatively, dsRNAEngineer also provides the design of a dsRNA fragment in targeting multiple pest genes by multi-target analysis (Figure 5B and C). Based on the user selected dsRNA fragments from different genes, a dsRNA containing all these fragments will be listed in the section of *dsRNA in targeting multiple genes* (Figure 5B). The organization order of these dsRNA fragments in a multi-target dsRNA will also be optimized by calculating the on- and off-targets using the same method described above (Figure 5C). The optimal dsRNA will be listed afterwards as the template for further dsRNA synthesis.

**Figure 5.**
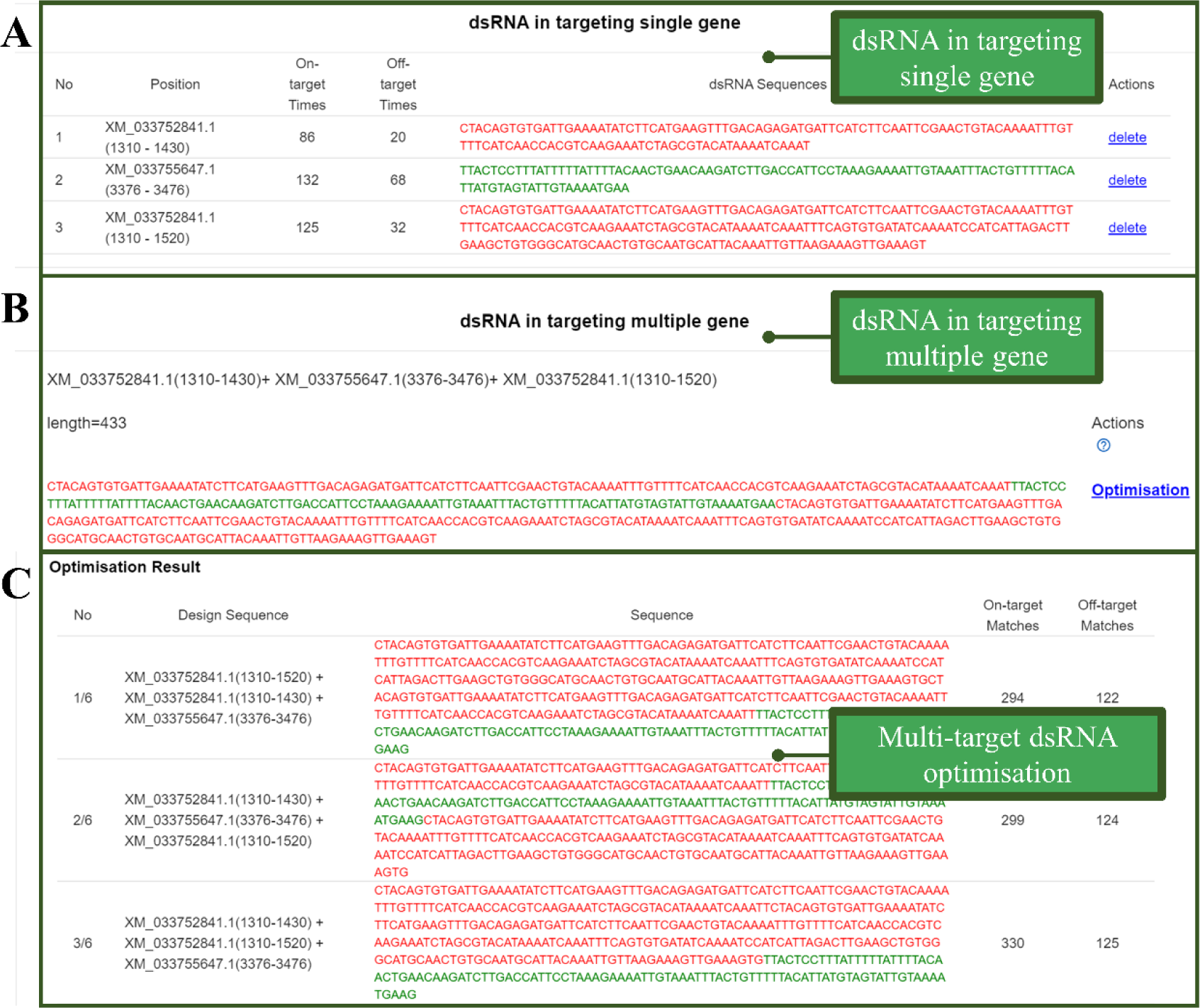
Multi-target analysis. (A) Post on-/off-target analysis, these dsRNA fragments targeting a single gene can be directly used as templates for further dsRNA synthesis. (B) These dsRNA fragments can also be optimized to target multiple pest genes by multi-target analysis. The fusion dsRNA sequences can then be taken as template for further dsRNA synthesis.

## Results

dsRNAEngineer is a comprehensive dsRNA design tool for pest control, including the species transcriptome-level analysis, on-target, and off-target. Additionally, we also provide screen-target for obtaining co-target genes per a group of pests, and multi-target analysis to optimize the organization order of different dsRNA fragments in a dsRNA on the basis of calculation of on- and off-targets. Both on- and off-targets used a same calculation principle, thus we performed study cases of screen-target and off-target analysis to validate the web server.

### Case 1: validation of screen-target analysis

For a group of closely related pests, the design of dsRNAs that target a gene (*β-Actin*) conserved in multiple spider mites (*T. urticae*, *T. evansi*, *T. cinnabarinus*, and *T. truncatus*) can achieve control of multiple pests with a conserved dsRNA fragment (24). By performing the screen-target analysis from dsRNAEngineer, there are 447 genes obtained with the cut-off value of sequence identity ≥ 97% & hit length ≥ 300bp. *β- Actin* ranked 123^rd^ in this analysis (Figure 6), demonstrating the usefulness of a tool like dsRNAEngineer in finding co-RNAi targets for various pest groups.

**Figure 6.**
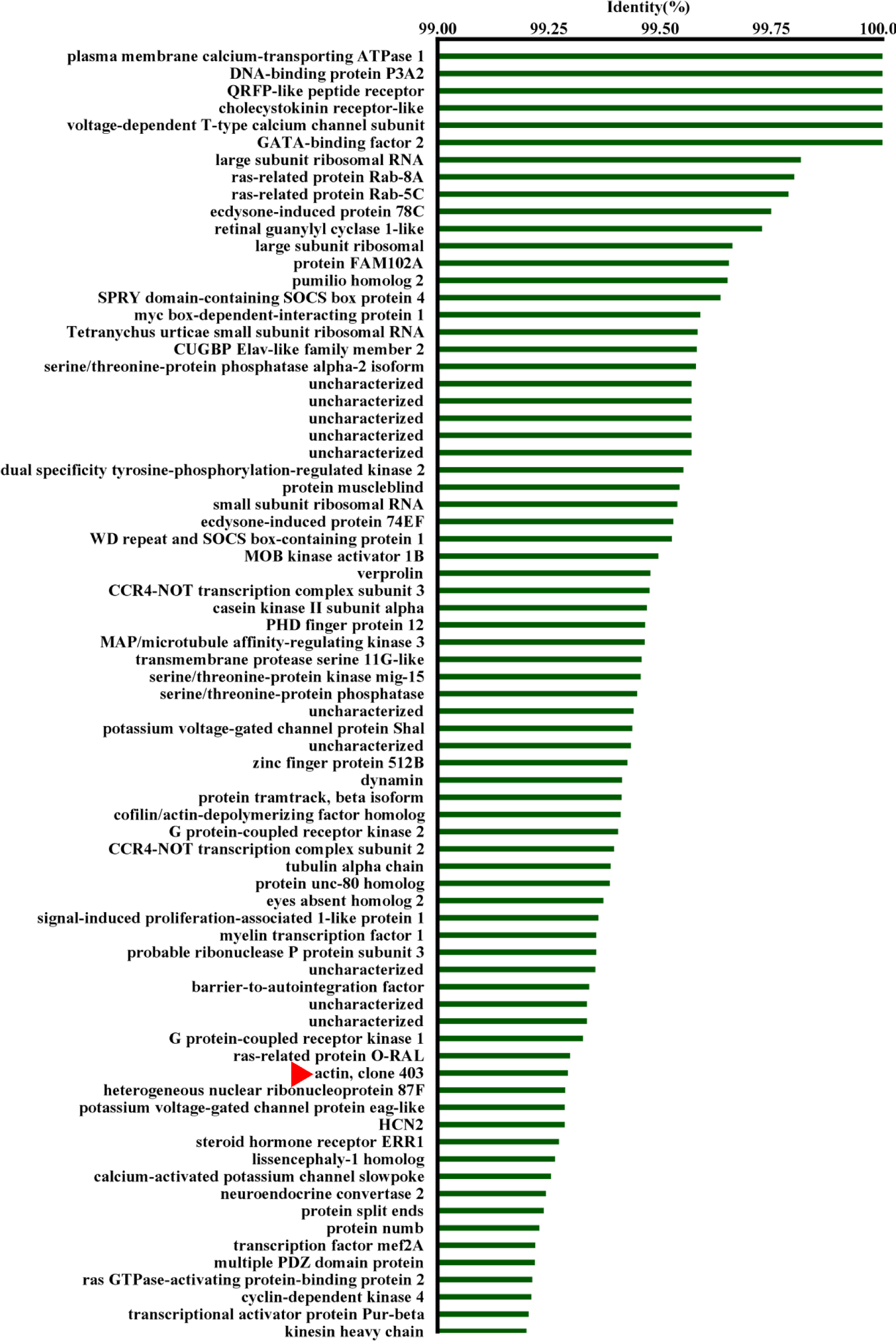
Screen-target analysis revealed candidate genes for co-RNAi targets to a group of spider mite pests. Key spider mite *Tetranychus urticae* was set as the major target pest, and three other spider mites including *T. evansi*, *T. cinnabarinus*, and *T. truncatus*, were selected co-targeted pests.

### Case 2: validation of off-target analysis

Two registered RNAi-based pest products (MON87411 and Ledprona) have been evaluated for their bio-safety, thus these dsRNA sequences are ideal cases to validate the off-target analysis. For this purpose, 84 non-pest species were selected for off-target analysis, in covering wide non-target-organism groups, such as aquatic ecosystems, birds, pollinators, silkworms, beneficial arthropods, earthworms, biocontrol fungi, crop plants, livestock animals, and mammals. Off-target analysis gave two datasets, transcriptome-level off-targets per an organism and the most off-target genes per an organism. For transcriptome-level off-target analysis per organism (Figure 8A and 8C for MON87411 and Ledprona, respectively), counts of off-targets are ranging from 2 to 134 for MON87411, and from 0 to 222 for Ledprona. For the most off-target genes per an organism (Figure 8B and 8D for MON87411 and Ledprona, respectively), counts of off-targets are ranging from 1 to 6 for MON87411, and from 0 to 22 for Ledprona. In summary, off-target analysis of MON87411 and Ledprona as positive controls suggest that dsRNAEngineer can be confidently used for dsRNA evaluation in pest control. It provides a reliable reference for assessing the safety of RNAi-based pest control.

**Figure 8.**
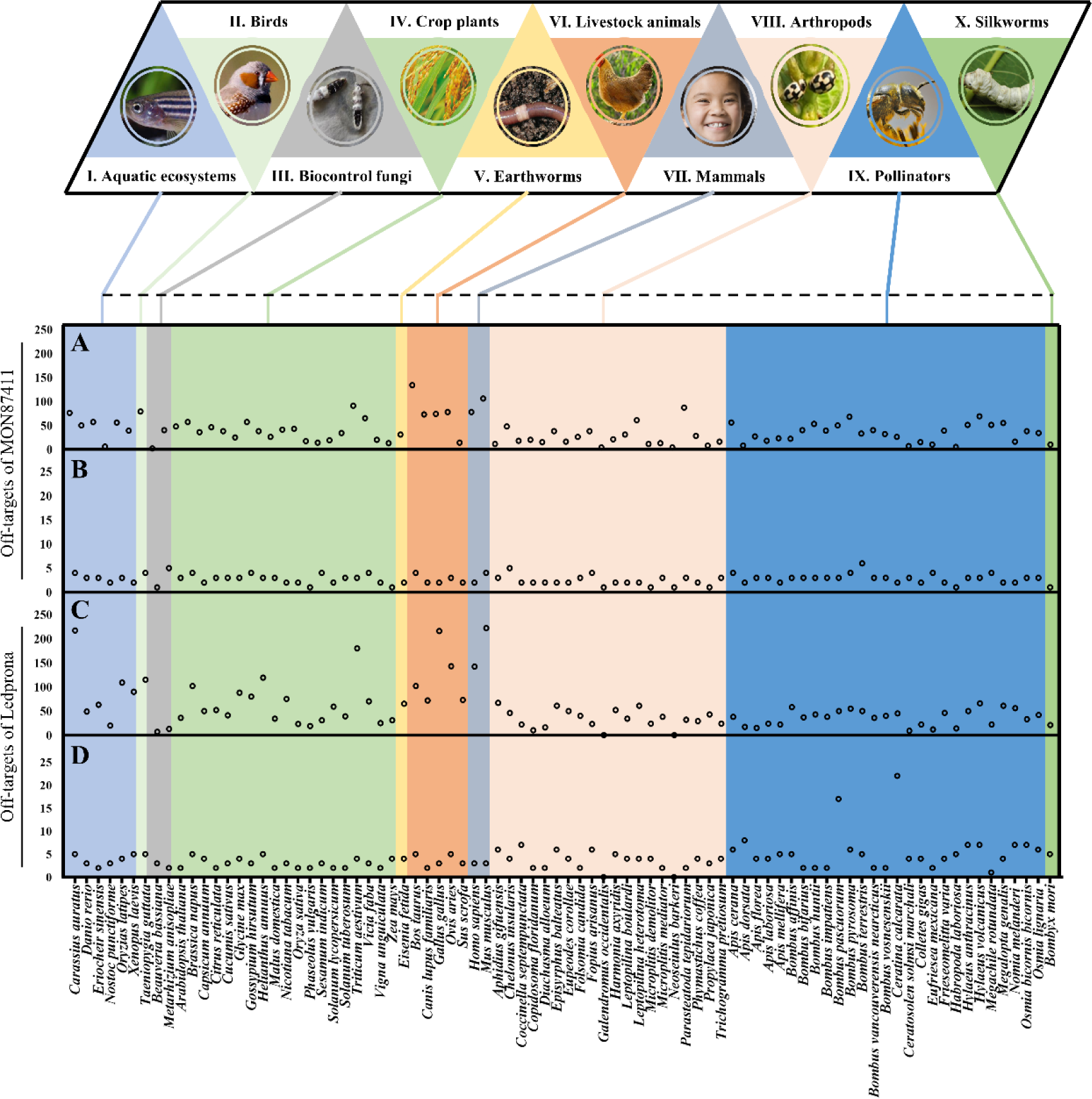
Validation of off-target analysis. (A) Counts of transcriptome-level off-targets per organism for MON87411 (dsSnf-7); (B) Counts of the most off-target genes per organism for MON87411 (dsSnf-7); (C) Counts of transcriptome-level off-targets per organism for Ledprona (dsV-ATP); (D) Counts of the most off-target genes per organism for Ledprona (dsV-ATP);

### Concluding remarks

In conclusion, dsRNAEngineer incorporates hundreds of pests and non-pests transcriptomes to allow a comprehensive dsRNA design for pest control, which enable users easily perform four functionalities including, screen-target, on-target, off-target, and multi-target to generate optimal dsRNA sequences for precise pest control. The online tool provides a user-friendly interface for this purpose. Limited species-selection or target gene input can initiate a task, and the result can be finalized in a relatively short time considering the transcriptome-level large data processing. Several cases including the commercialized dsRNA tested above, demonstrate that dsRNAEngineer is a powerful tool for dsRNA design. We believe this tool will not only reshape the current concept of dsRNA design for pest control, but also the efficient way to evaluate the biosafety of insecticidal dsRNA.

## Data availability

dsRNAEngineer is freely available at https://dsrna-engineer.cn/. It is free and open to all users without login requirement.

## Supplmentary Figures

**Supplemental Figure 1.**
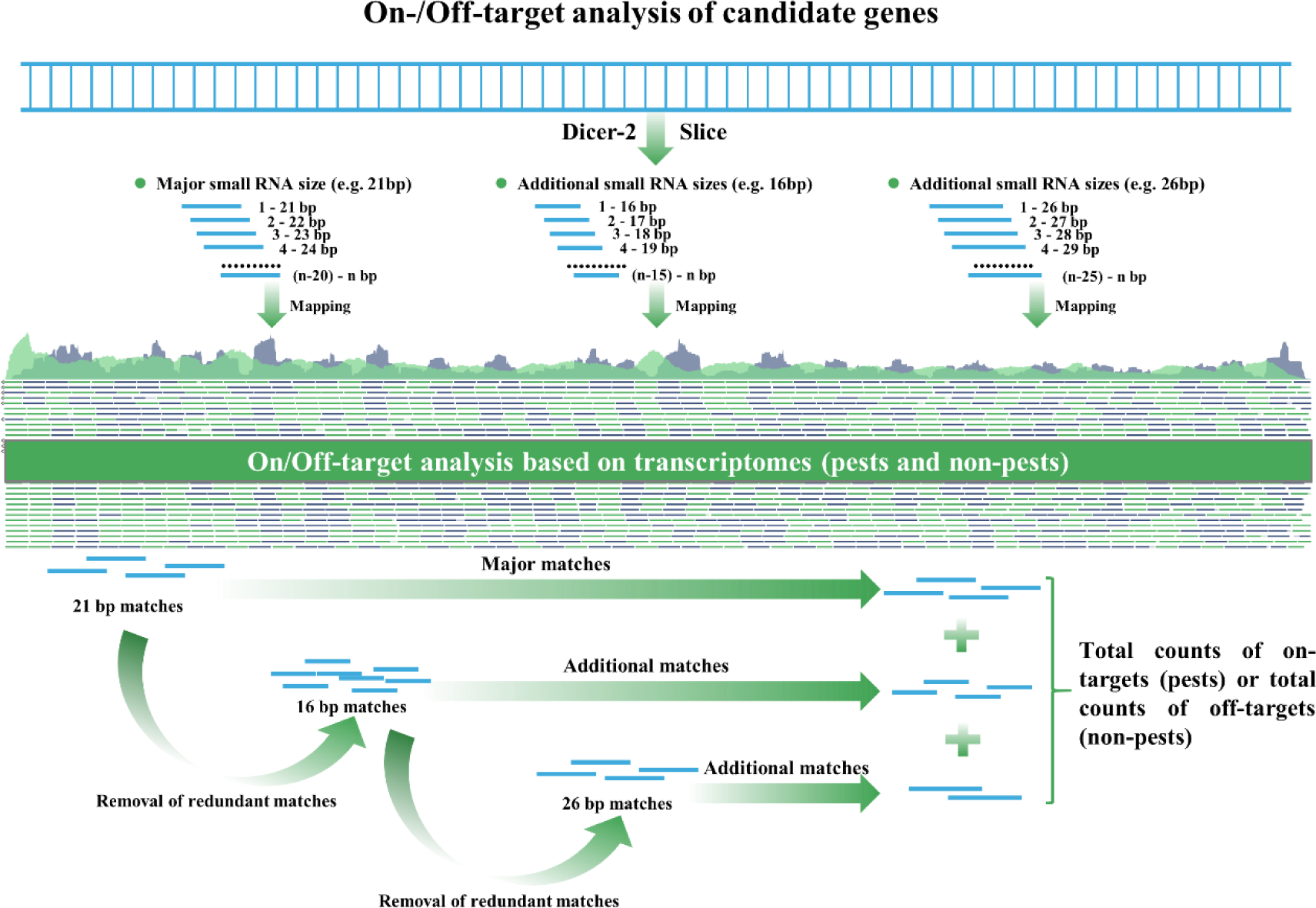
On-/Off-target analysis of candidate genes on the dsRNAEngineer. dsRNAEngineer slices the candidate target genes into major small RNA size (e.g. 21bp) and two additional small RNA sizes (e.g. 16bp and 26bp), then performs On/Off-target analysis based on transcriptomes, and finally counts the total on-targets and off-targets for candidate target genes.

**Supplemental Figure 2.**
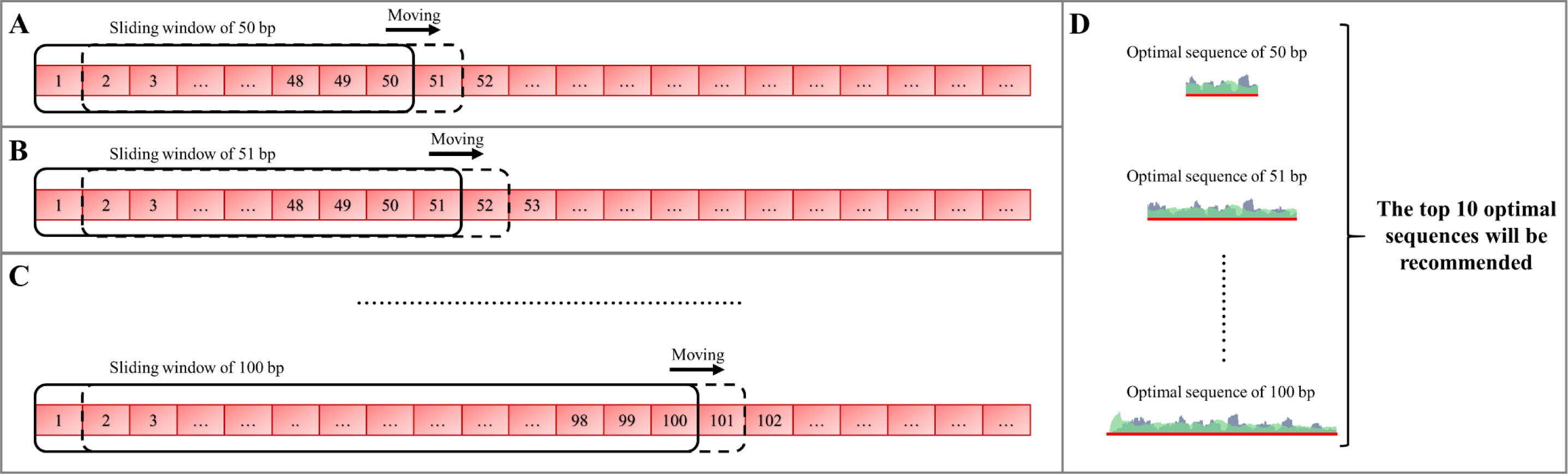
Sliding window analysis of 50-100bp dsRNA fragments on the dsRNAEngineer platform. (A) the start length of the sliding window analysis is 50bp; (B-C) After one round of analysis covering the whole target gene sequence, 50+1 bp shift is taken as the next sliding window analysis until 100 bp is reached; (D) Each sliding window will generate an optimal sequence, and all optimal sequences from each sliding window will be re-ranked and top ten will be recommended.

## References

1. Fire, A., Xu, S.Q., Montgomery, M.K., Kostas, S.A., Driver, S.E. and Mello, C.C. (1998) Potent and specific genetic interference by double-stranded RNA in *Caenorhabditis elegans*. Nature, 391, 806–811.

2. Pretty, J., Benton, T.G., Bharucha, Z.P., Dicks, L.V., Flora, C.B., Godfray, H.C.J., Goulson, D., Hartley, S., Lampkin, N., Morris, C. et al. (2018) Global assessment of agricultural system redesign for sustainable intensification. Nat. Sustain., 1, 441–446.

3. Tang, F.H.M., Lenzen, M., McBratney, A. and Maggi, F. (2021) Risk of pesticide pollution at the global scale. Nat. Geosci., 14, 206–210.

4. Belles, X. (2010) Beyond *Drosophila*: RNAi In Vivo and Functional Genomics in Insects. Annu. Rev. Entomol., 55, 111–128.

5. Zhu, K.Y. and Palli, S.R. (2020) Mechanisms, Applications, and Challenges of Insect RNA Interference. Annu. Rev. Entomol., 65, 293–311.

6. Mezzetti, B., Smagghe, G., Arpaia, S., Christiaens, O., Dietz-Pfeilstetter, A., Jones, H., Kostov, K., Sabbadini, S., Opsahl-Sorteberg, H., Ventura, V. et al. (2020) RNAi: What is its position in agriculture? J. Pest Sci., 93, 1125–1130.

7. Niu, J., Chen, R. and Wang, J. (2024) RNA interference in insects: the link between antiviral defense and pest control. Insect Sci., 31, 2–12.

8. Naegeli, H., Birch, A.N., Casacuberta, J., De Schrijver, A., Gralak, M.A., Guerche, P., Jones, H., Manachini, B., Messean, A., Nielsen, E.E., et al. (2018) Assessment of genetically modified maize MON 87411 for food and feed uses, import and processing, under Regulation (EC) No 1829/2003 (application EFSA-GMO-NL-2015-124). EFSA J., 16, e05310.

9. Yan, J., Nauen, R., Reitz, S., Alyokhin, A., Zhang, J., Mota-Sanchez, D., Kim, Y., Palli, S.R., Rondon, S.I., Nault, B.A. et al. (2024) The new kid on the block in insect pest management: sprayable RNAi goes commercial. Sci. China Life Sci. doi: 10.1007/s11427-024-2612-1.

10. Xie, Z.X., Allen, E., Wilken, A. and Carrington, J.C. (2005) DICER-LIKE 4 functions in trans-acting small interfering RNA biogenesis and vegetative phase change in *Arabidopsis thaliana*. Proc. Natl. Acad. Sci. U. S. A., 102, 12984–12989.

11. Flemr, M., Malik, R., Franke, V., Nejepinska, J., Sedlacek, R., Vlahovicek, K. and Svoboda, P. (2013) A Retrotransposon-Driven Dicer Isoform Directs Endogenous Small Interfering RNA Production in Mouse Oocytes. Cell, 155, 807–816.

12. Chen, J., Peng, Y., Zhang, H., Wang, K., Zhao, C., Zhu, G., Reddy Palli, S. and Han, Z. (2021) Off- target effects of RNAi correlate with the mismatch rate between dsRNA and non-target mRNA. RNA Biol., 18, 1747–1759.

13. Chen, J., Sheng, C., Peng, Y., Wang, K., Jiao, Y., Palli, S.R. and Cao, H. (2023) Transcript Level and Sequence Matching Are Key Determinants of Off-Target Effects in RNAi. J. Agric. Food Chem., 72, 577–589.

14. Wang, Z., Chen, R., Jiang, Y., Wang, Z., Wang, J. and Niu, J. (2023) Investigation of potential non- target effects to a ladybeetle *Propylea japonica* in the scenario of RNAi-based pea aphid control. Entomol. Gen., 43, 79–88.

15. Ma, W., Wu, T., Zhang, Z., Li, H., Situ, G., Yin, C., Ye, X., Chen, M., Zhao, X., He, K. et al. (2022) Using transcriptome Shannon entropy to evaluate the off-target effects and safety of insecticidal siRNAs. J. Integr. Agric., 21, 170–177.

16. Booker, M., Samsonova, A.A., Kwon, Y., Flockhart, I., Mohr, S.E. and Perrimon, N. (2011) False negative rates in *Drosophila* cell-based RNAi screens: a case study. BMC Genomics, 12, 50.

17. Henschel, A., Buchholz, F. and Habermann, B. (2004) DEQOR: a web-based tool for the design and quality control of siRNAs. Nucleic Acids Res., 32, W113–W120.

18. Naito, Y., Yamada, T., Matsumiya, T., Ui-Tei, K., Saigo, K. and Morishita, S. (2005) dsCheck: highly sensitive off-target search software for double-stranded RNA-mediated RNA interference. Nucleic Acids Res., 33, W589–W591.

19. Horn, T. and Boutros, M. (2010) E-RNAi: a web application for the multi-species design of RNAi reagents-2010 update. Nucleic Acids Res., 38, W332–W339.

20. Chen, S., Zhou, Y., Chen, Y. and Gu, J. (2018) fastp: an ultra-fast all-in-one FASTQ preprocessor. Bioinformatics, 34, 884–890.

21. Grabherr, M.G., Haas, B.J., Yassour, M., Levin, J.Z., Thompson, D.A., Amit, I., Adiconis, X., Fan, L., Raychowdhury, R., Zeng, Q. et al. (2011) Full-length transcriptome assembly from RNA-Seq data without a reference genome. Nat. Biotechnol., 29, 644–U130.

22. Altschul, S.F., Gish, W., Miller, W., Myers, E.W. and Lipman, D.J. (1990) BASIC LOCAL ALIGNMENT SEARCH TOOL. J. Mol. Biol., 215, 403–410.

23. Shang, F., Ding, B., Ye, C., Yang, L., Chang, T., Xie, J., Tang, L., Niu, J. and Wang, J. (2020) Evaluation of a cuticle protein gene as a potential RNAi target in aphids. Pest Manag. Sci., 76, 134–140.

24. Wu, M., Zhang, Q., Dong, Y., Wang, Z., Zhan, W., Ke, Z., Li, S., He, L., Ruf, S., Bock, R. et al. (2023) Transplastomic tomatoes expressing double-stranded RNA against a conserved gene are efficiently protected from multiple spider mites. New Phytol., 237, 1363–1373.

25. Dong, Y., Zhang, Q., Mao, Y., Wu, M., Wang, Z., Chang, L. and Zhang, J. (2024) Control of two insect pests by expression of a mismatch corrected double-stranded RNA in plants. Plant Biotechnol. J., 22, 2010–2019.

